# Genetic Prediction of Male Pattern Baldness

**DOI:** 10.1101/072306

**Authors:** Saskia P Hagenaars, W David Hill, Sarah E Harris, Stuart J Ritchie, Gail Davies, David C Liewald, Catharine R Gale, David J Porteous, Ian J Deary, Riccardo E Marioni

**Affiliations:** Centre for Cognitive Ageing and Cognitive Epidemiology, University of Edinburgh, Edinburgh, UK; Department of Psychology, University of Edinburgh, Edinburgh, UK; Division of Psychiatry, University of Edinburgh, Edinburgh, UK; Medical Genetics Section, Centre for Genomic and Experimental Medicine, Institute of Genetics and Molecular Medicine, University of Edinburgh, Edinburgh, UK; Medical Research Council Lifecourse Epidemiology Unit, University of Southampton, Southampton, UK; Queensland Brain Institute, The University of Queensland, Brisbane, QLD, Australia

## Abstract

Male pattern baldness can have substantial psychosocial effects, and it has been phenotypically linked to adverse health outcomes such as prostate cancer and cardiovascular disease. We explored the genetic architecture of the trait using data from over 52,000 male participants of UK Biobank, aged 40-69 years. We identified over 250 independent novel genetic loci associated with severe hair loss. By developing a prediction algorithm based entirely on common genetic variants, and applying it to an independent sample, we could discriminate accurately (AUC = 0.82) between those with no hair loss from those with severe hair loss. The results of this study might help identify those at the greatest risk of hair loss and also potential genetic targets for intervention.

## Introduction

Male pattern baldness affects around 80% of men by the age of 80 years [1] and it can have substantial psychosocial impacts via changes in self-consciousness and social perceptions [2, 3]. In addition to alterations in physical appearance, some, but not all, studies have identified negative health outcomes associated with baldness including increased risk of prostate cancer [4-6] and cardiovascular disease [7-9]. Baldness is known to be substantially heritable [10]. Here, we use a large population-based dataset to identify many of the genes linked to variation in baldness, and build a genetic score to improve prediction of severe hair loss.

The total proportion of variance in male pattern baldness that can be attributed to genetic factors has been estimated in twin studies to be approximately 80% for both early- and late-onset hair loss [11, 12]. The remaining variance in these twin studies was attributable to non-shared environmental factors. Newer molecular-genetic methods have estimated the autosomal single-nucleotide polymorphism (SNP)-based, common-variant heritability of baldness at around 50% [13]. Molecular methods also indicate some overlap between genetic variants linked to baldness and those linked to phenotypes such as height, waist-hip ratio, age at voice drop in males, age at menarche in females, and presence of a unibrow [14].

A number of studies have identified specific genetic variants linked to variations in baldness, usually with the *AR* gene showing the strongest association. The largest published genome-wide association study (GWAS) to date highlighted eight independent genetic loci that were linked to baldness; the top *AR* SNP yielded an odds ratio of 2.2 in a case-control meta-analysis of 12,806 individuals of European ancestry [15]. One of the autosomal hits identified in that study was found to be in a gene linked to Parkinson’s disease. More recently, a review paper highlighted fifteen loci from six studies that have been associated at genome-wide significance (P<5×10^−8^) with baldness; two of the loci were located on the X chromosome [16].

Several attempts have been made to build predictors of male pattern baldness using polygenic risk scores. Heilman et al. found, using a case-control design with ∼600 per arm, that a predictor based on 34,186 SNPs explained 4.5% of the variance on the liability scale [17]. Marcinska et al. used candidate genes to build 5-SNP and 20-SNP polygenic predictors, which performed quite well when considering prediction of early-onset male pattern baldness, but poorly when considering those with no baldness versus those with severe baldness across all ages [18]. Most recently, a 20-SNP predictor was assessed in three European studies [13]. It achieved a maximum Area Under the Curve (AUC) prediction of 0.74 in an early-onset cohort, but weaker estimates in the other two, late-onset cohorts (AUC = 0.69 and 0.71). These values correspond to poor-to-fair predictions of baldness. In that study, age was included in the predictor, explaining the bulk of the differences. A meta-analysis of the three cohorts’ GWAS studies identified a novel locus on chromosome 6. The study also estimated the SNP-based heritability of early-onset (56% (SE 22%) from the autosomes, 23% (SE 1.1%) from the X chromosome) and late-onset baldness (42% (SE 23%) from the autosomes, 10% (SE 5%) from the X chromosome).

### The Present Study

The UK Biobank study [19] (http://www.ukbiobank.ac.uk) is a large, population-based genetic epidemiology cohort. At its baseline assessment (2006-2010), around 500,000 individuals aged between 40 and 70 years and living in the UK completed health and lifestyle questionnaires and provided biological samples for research.

The present study reports a GWAS of male pattern baldness in the UK Biobank cohort. It is over four times the size of the previously-largest meta-analytic study [15]. We then split the cohort into a large ‘discovery’ sample and smaller ‘test’ sample to perform a prediction analysis, determining the accuracy of a polygenic profile score built from the GWAS findings in discriminating between those with severe hair loss and those with no hair loss.

## Results

The mean age of the 52,874 men was 57.2 years (SD 8.0). 16,724 (31.6%) reported no hair loss, 12,135 (23.0%) had slight hair loss, 14,234 (26.9%) had moderate hair loss, and 9,781 (18.5%) had severe hair loss.

The genome-wide association study of the four-category self-reported baldness measure in 52,874 White British men from UK Biobank yielded 13,029 autosomal hits from the imputed data (P<5×10^−8^) in addition to 117 hits (out of 14,350 genotyped SNPs) on the X chromosome (**Figure 1**). The QQ plot for the autosomal GWAS is shown in **Figure S1**. An LD clumping analysis indicated that these hits can be attributed to 247 independent autosomal regions. All of the 8 autosomal hits identified by Li et al. [15] and the additional hits noted in Heilmann-Heimbach et al. [16] replicated in the UK Biobank dataset with a maximum P value of 6.4×10^−14^ (**Table 1**). The previously reported X chromosome variant from Li et al. [15] and the variant from Richards et al. [10] also replicated with P-values that were effectively zero (**Table 1**). The chromosome 6 hit from Liu et al. [13] failed to replicate (P=0.37). A list of the top 20 independent autosomal hits is presented in **Table 2**. The top 10 independent X chromosome hits are presented in **Supplementary Table 1**; rs140488081 and rs7061504 are intronic SNPs in the *OPHN1* gene.

**Figure 1.**
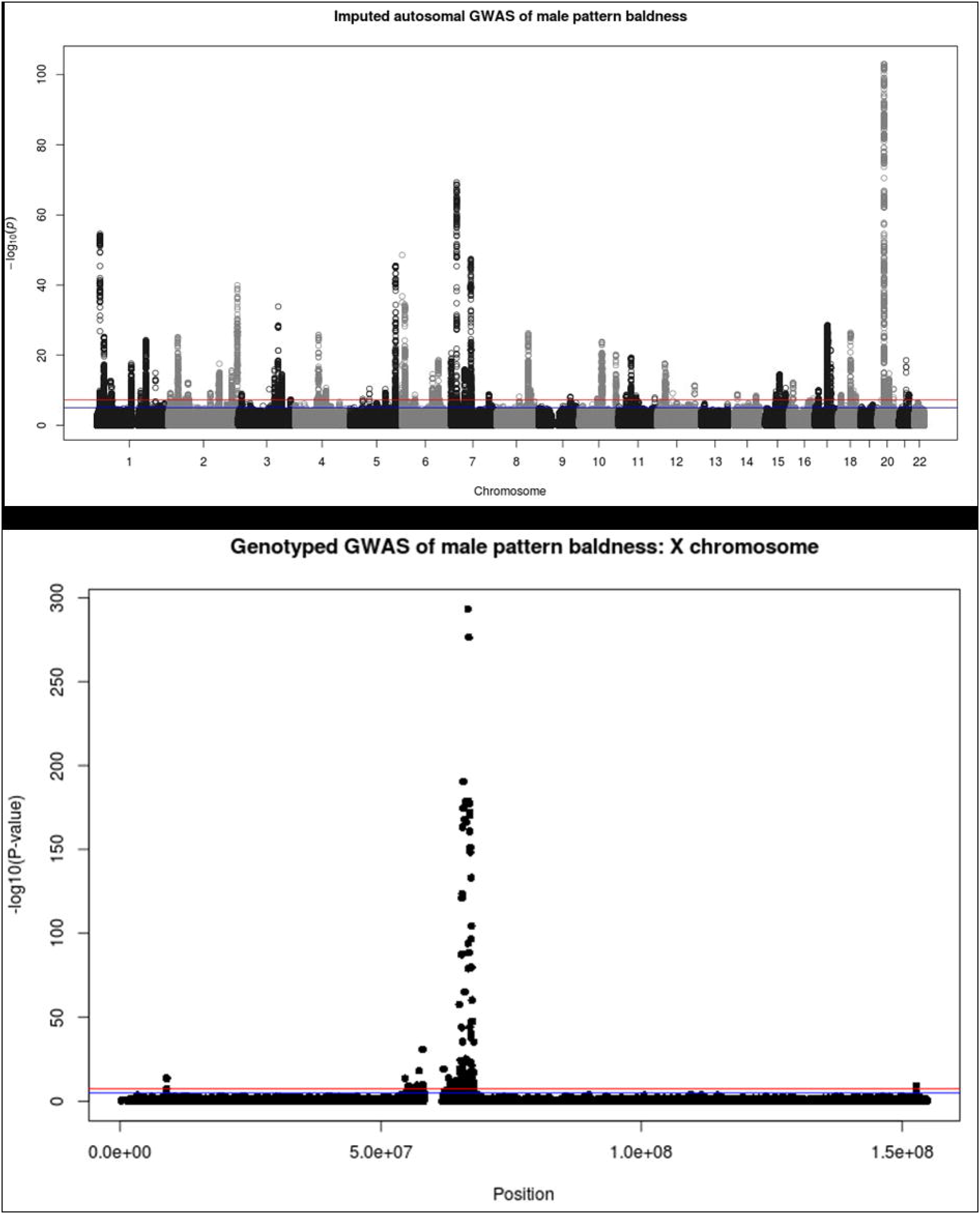
Manhattan Plots of imputed autosomal GWAS (top panel) and genotyped X chromosome GWAS (bottom panel) of male pattern baldness.

**Table 1.**
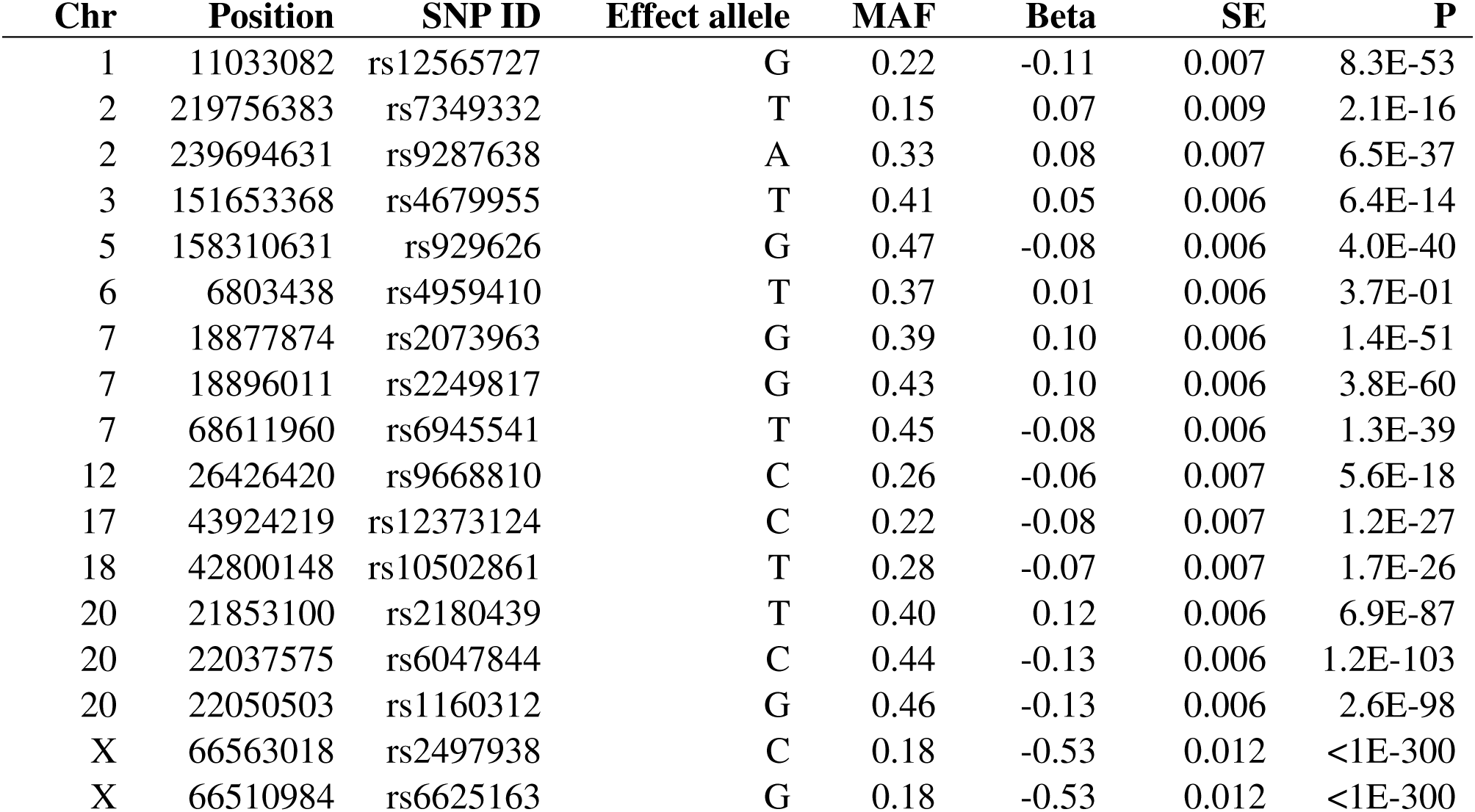
Lookup of GWAS hits from Li et al. 2012 [15], Heilmann-Heimbach et al. 2016 [16] and Liu et al. 2016 [13].

| Chr | Position | SNP ID | Effect allele | MAF | Beta | SE | P |
| --- | --- | --- | --- | --- | --- | --- | --- |
| 1 | 11033082 | rs12565727 | G | 0.22 | -0.11 | 0.007 | 8.3E-53 |
| 2 | 219756383 | rs7349332 | T | 0.15 | 0.07 | 0.009 | 2.1E-16 |
| 2 | 239694631 | rs9287638 | A | 0.33 | 0.08 | 0.007 | 6.5E-37 |
| 3 | 151653368 | rs4679955 | T | 0.41 | 0.05 | 0.006 | 6.4E-14 |
| 5 | 158310631 | rs929626 | G | 0.47 | -0.08 | 0.006 | 4.0E-40 |
| 6 | 6803438 | rs4959410 | T | 0.37 | 0.01 | 0.006 | 3.7E-01 |
| 7 | 18877874 | rs2073963 | G | 0.39 | 0.10 | 0.006 | 1.4E-51 |
| 7 | 18896011 | rs2249817 | G | 0.43 | 0.10 | 0.006 | 3.8E-60 |
| 7 | 68611960 | rs6945541 | T | 0.45 | -0.08 | 0.006 | 1.3E-39 |
| 12 | 26426420 | rs9668810 | C | 0.26 | -0.06 | 0.007 | 5.6E-18 |
| 17 | 43924219 | rs12373124 | C | 0.22 | -0.08 | 0.007 | 1.2E-27 |
| 18 | 42800148 | rs10502861 | T | 0.28 | -0.07 | 0.007 | 1.7E-26 |
| 20 | 21853100 | rs2180439 | T | 0.40 | 0.12 | 0.006 | 6.9E-87 |
| 20 | 22037575 | rs6047844 | C | 0.44 | -0.13 | 0.006 | 1.2E-103 |
| 20 | 22050503 | rs1160312 | G | 0.46 | -0.13 | 0.006 | 2.6E-98 |
| X | 66563018 | rs2497938 | C | 0.18 | -0.53 | 0.012 | <1E-300 |
| X | 66510984 | rs6625163 | G | 0.18 | -0.53 | 0.012 | <1E-300 |

**Table 2.**
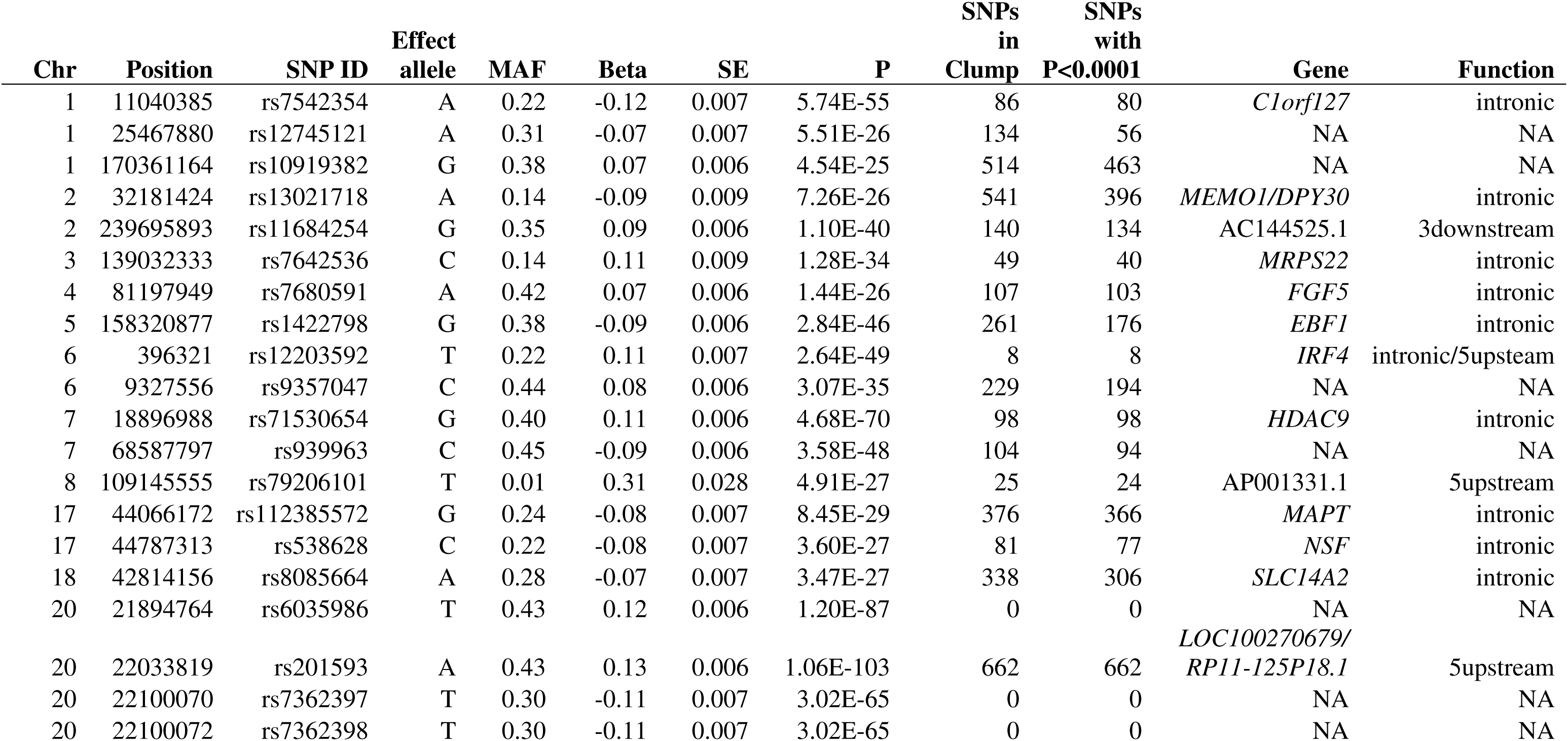
Top 20 independent autosomal GWAS hits

The gene-based analysis identified 112 autosomal genes and 13 X chromosome genes that were associated with baldness after a Bonferroni correction (P<0.05/18,061 and P<0.05/567, respectively). The top gene-based hit was, as expected, the androgen receptor on the X chromosome (P=2.0×10^−269^). A full list of the autosomal significant gene-based hits is provided in **Supplementary Table 2** and significant genes on the X chromosome are shown in **Supplementary Table 3**.

Using common genetic variants with a minor allele frequency of at least 1%, GCTA-GREML analysis found that 47.3% (SE 1.3%) of the variance in baldness can be explained by common autosomal genetic variants, 4.6% (SE 0.3%) can be explained by common X chromosome variants.

Genetic correlations were examined between male pattern baldness and numerous (n=22) cognitive, health, and anthropometric traits using LD Score regression. No significant associations were found and all estimates were close to the null (**Supplementary Table 4**). The GWAS for self-reported baldness was re-run on a sub-sample of 40,000 individuals - retaining an equal proportion of each of the four baldness patterns as observed in the full cohort - to allow a polygenic prediction score to be built and applied to the remaining, independent sample of 12,874 individuals. The most powerful predictions from comparing the extreme phenotype groups were observed at the P<1×10^−5^ threshold for the autosomal polygenic score and at P<1 for the X chromosome polygenic score (**Table 3**). The optimal autosomal polygenic score yielded an AUC of 0.75 for discriminating between those with no hair loss (n=4,123) and those with severe hair loss (n=2,456). The AUC values between those with no hair loss and those with slight and moderate hair loss were 0.586 and 0.652, respectively. The corresponding AUCs for the X chromosome polygenic score were 0.724 (severe loss), 0.592 (moderate loss), and 0.523 (slight loss, P=0.001), when examined against the reference group of no hair loss. All logistic regression P-values for the effect of the polygenic score were <2×10^−16^ except where noted. An additive combination of the autosomal and X chromosome polygenic scores gave an AUC of 0.816 for severe baldness, 0.707 for moderate baldness, and 0.619 for slight baldness (**Figure 2**).

**Figure 2.**
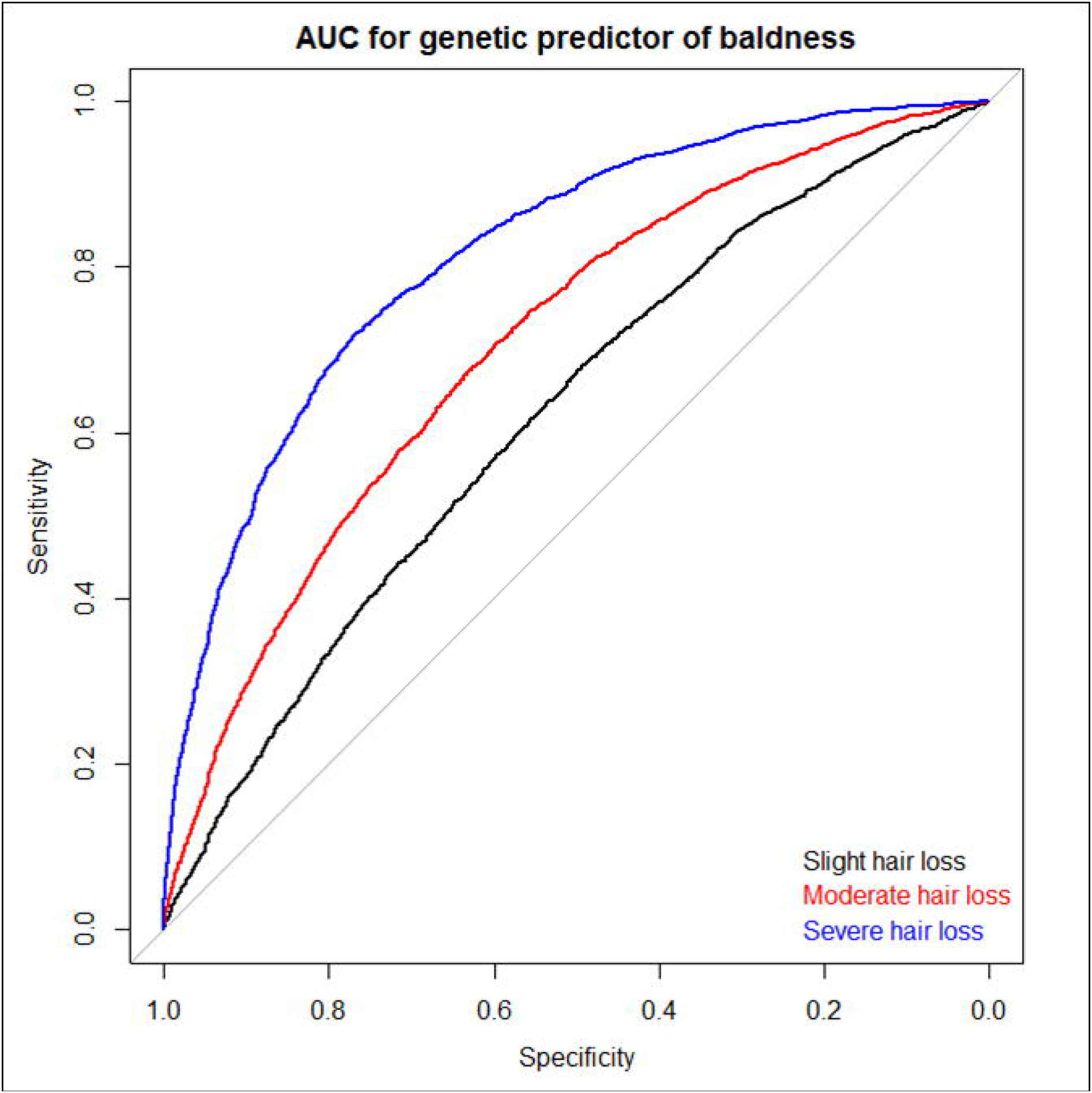
Area under the curve plot for discriminating those with hair loss from those with no loss.

**Table 3.** AUC results for severe hair loss versus no hair loss for all autosomal and X chromosome polygenic thresholds.

| **Genomic region** | **P value threshold** | **n SNPs** | **AUC** |
| --- | --- | --- | --- |
| Autosomal | <1x10-20 | 37 | 0.648 |
|  | <5x10-8 | 224 | 0.725 |
|  | <1x10-5 | 546 | 0.748 |
|  | <0.01 | 10581 | 0.725 |
|  | <0.05 | 40047 | 0.701 |
|  | <0.10 | 74068 | 0.687 |
|  | <0.50 | 322885 | 0.663 |
|  | <1 | 622739 | 0.662 |
| X chromosome | <1x10-20 | 52 | 0.611 |
|  | <5x10-8 | 109 | 0.630 |
|  | <1x10-5 | 159 | 0.625 |
|  | <0.01 | 479 | 0.649 |
|  | <0.05 | 1139 | 0.673 |
|  | <0.10 | 1914 | 0.691 |
|  | <0.50 | 7761 | 0.720 |
|  | <1 | 14653 | 0.724 |

The results of the partitioned heritability analysis indicated that 27 of the functional annotations from the baseline model were statistically significant (**Supplementary Figure 2** and **Supplementary Table 5**). These significant annotations included a broad array of functional elements including histone marks, enhancer regions, conserved regions, and DNaseI hypersensitivity sites (DHS). The ten tissue types were then tested for significance after controlling for the baseline model. Following correction for multiple testing all ten of the tissue groups showed significant enrichment (**Supplementary Figure 3** and **Supplementary Table 5**).

## Discussion

In this large GWAS study of male pattern baldness, we identified 287 independent genetic signals that are linked to differences in the trait, a substantial advance over the previous largest GWAS meta-analysis, which identified eight independent signals [15]. We showed, in line with a previous study [13], but with much greater precision, that a substantial proportion of individual differences in hair loss patterns can be explained by common genetic variants on the autosomes as well as the X chromosome. Finally, by splitting our cohort into a discovery and a prediction sample, we show a good predictive discrimination (AUC = 0.82) between those with no hair loss and those with severe hair loss.

Despite there being genetic overlap for SNP hits associated with baldness and Parkinson’s Disease—first noted in Li et al. [15] and replicated here—we observed no genetic correlations between baldness and any of the health, cognitive, or anthropometric outcomes we studied. The point estimates for the genetic correlations were all near zero, suggesting true null associations as opposed to a lack of statistical power to detect modest-sized correlations.

As mentioned above, the GWAS identified 247 independent autosomal loci and 40 independent X chromosome loci. The top 20 hits from the autosomes were located in/near to genes that have been associated with, for example, hair growth/length in mice (*FGF5*) [20], grey hair (*IRF4*) [21], cancer (breast: *MEMO1* [22], bladder: *SLC14A2* [23]), histone acetylation (*HDAC9*), and frontotemporal dementia (*MAPT*) [24]. Two of the top 10 X chromosome SNPs were located in *OPHN1,* a gene previously associated with X-linked mental retardation [25].

Of the top autosomal gene-based findings (maximum P=3.1×10^−15^), *RSPO2* has been linked to hair growth in dogs. *PGDFA* has been linked to hair follicle development [26]; *EBF1* is expressed in dermal papillae in mature hair follicles [27]; *PRR23B* is proximal to a GWAS hit for eyebrow thickness [21]; and *WNT10A* has been linked to both straight hair [28] and dry hair [29].

The top X chromosome gene-based findings included the androgen receptor (*AR*), which has been well established as a baldness associated gene [30], along with its upstream (*EDA2R*) and downstream (*OPHN1*) genes. *EDA2R* plays a role in the maintenance of hair and teeth as part of the tumor necrosis factor receptor. Onset of male pattern baldness could by influenced by *EDA2R* via activation of nuclear proto-oncoprotein *c-Jun*, which is linked to transcription activation of *AR* [31]. Two other genes included in the gene-based findings, *OPHN1* and *ZC4H2*, have previously been associated with X-linked mental retardation [25, 32].

The optimal thresholds from the prediction results highlight the polygenicity of the baldness trait and suggest that there may be many opportunities to identify novel pathways and possible therapies linked to baldness. The release of the full set of UK Biobank genotypes at the beginning of the next calendar year (2017) will likely enhance the predictive power of the polygenic score, and will probably allow for the identification of even more novel genetic loci in future GWAS studies.

The main strength of this study is its large sample size and phenotypic homogeneity. Many meta-analytic studies of complex traits are weakened by different cohorts collecting data at different time-points, under different protocols, in different populations. The present study replicated all of the previously identified autosomal hits for baldness from Li et al.[15] and Heilmann-Heimbach et al., [16] suggesting a degree of robustness in phenotypic measurement. Whereas the genomic inflation factor from the GWAS was large (1.09, Q-Q plot in **Supplementary Figure 1**), this is likely to be a result of genuine polygenic effects. We have used identical analysis protocols for other traits with far lower SNP-based heritabilities in the same UK Biobank cohort and observed no evidence of inflation [33].

### Conclusion

We identified hundreds of independent, novel genetic correlates of male pattern baldness - an order of magnitude greater than the list of previous genome-wide hits. Our top SNP and gene-based hits were in genes that have previously been associated with hair growth and development. We also generated a powerful polygenic predictor that discriminated well between those with no hair loss and those with severe hair loss.

## Methods

### Data

Data came from the first release of genetic data of the UK Biobank study and analyses were performed under the data application 10279. Ethical approval for UK Biobank was granted by the Research Ethics Committee (11/NW/0382).

Genotyping details including quality control steps have been reported previously [33]. Briefly, from the sample with genetic data available as of June 2015, 112,151 participants remained after the following exclusion criteria were applied: SNP missingness, relatedness, gender mismatch, non-British ancestry, and failed quality control for the UK BiLEVE study [33]. For the current analysis, an imputed dataset was used for the autosomes (reference set panel combination of the UK10K haplotype and 1000 Genomes Phase 3 panels: http://biobank.ctsu.ox.ac.uk/crystal/refer.cgi?id=157020). Imputed data were not available for the X chromosome; hence only genotyped variants were considered. X chromosome quality control steps included a minor allele frequency cut-off of 1% and a genotyping call rate cutoff of 98% [34]. For the imputed autosomal data, we restricted the analyses to variants with a minor allele frequency >0.1% and an imputation quality score >0.1.

From the sample of 112,151 unrelated White British participants with genetic data, we identified 52,874 men with a self-reported response to UK Biobank question 2395, which was adapted from the Hamilton-Norwood scale [35, 36]. These men were asked to choose, from four patterns (no loss; slight loss; moderate loss; severe loss), the one that matched their hair coverage most closely. **Figure 3** shows a screenshot of the four options.

**Figure 3.**
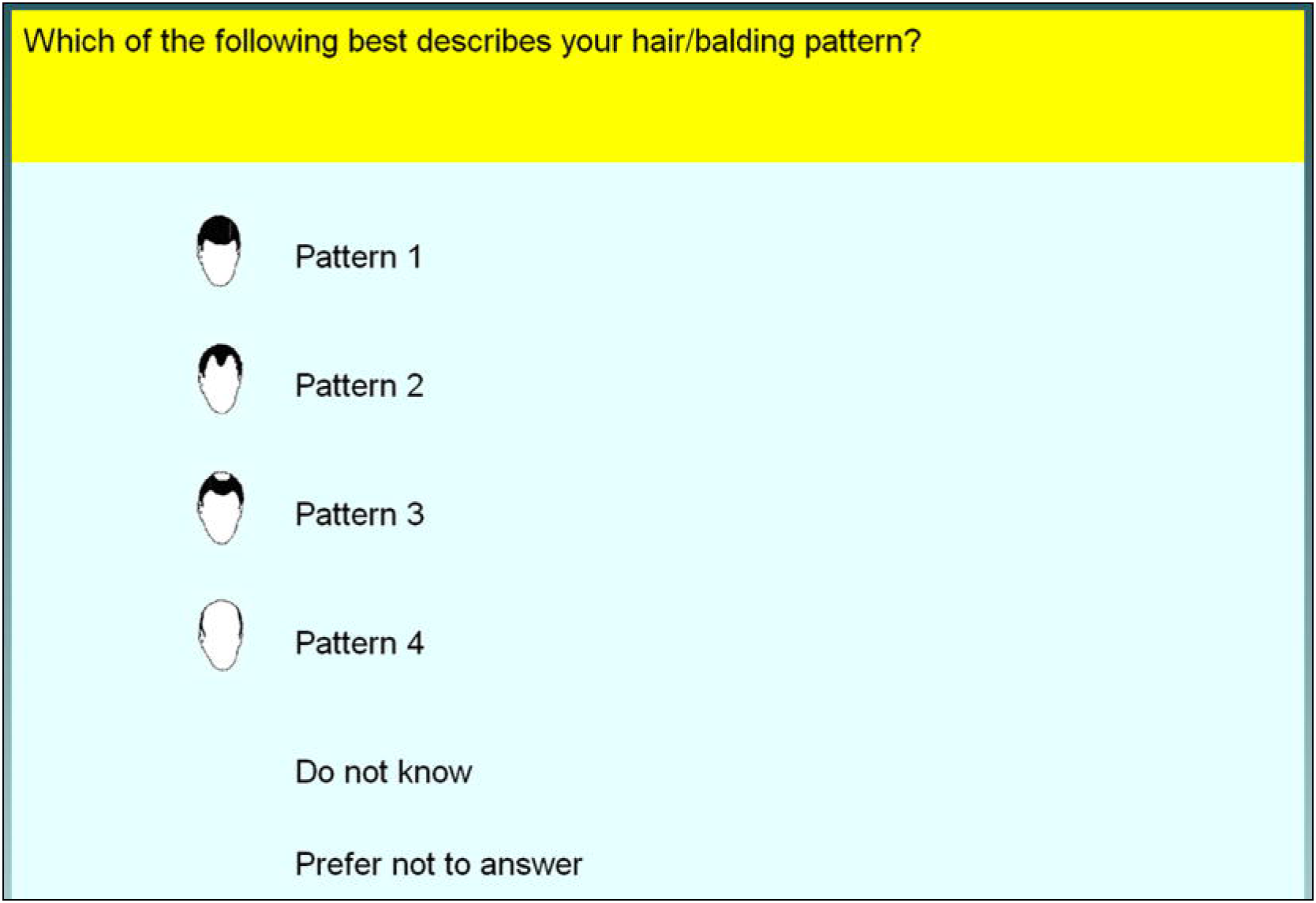
Screenshot of UK Biobank question 2395 on male-pattern baldness

### Statistical Analysis

A genome-wide association study was conducted using baldness pattern residuals as the dependent variable. The residuals were obtained from a linear regression model of baldness pattern on age, assessment centre, genotyping batch and array, and 10 principal components to correct for population stratification.

The GWAS for the imputed autosomal dataset was performed in SNPTest v2.5.1 [37] via an additive model, using genotype probability scores. The GWAS for the X chromosome was performed in PLINK [38, 39].

The number of independent signals from the GWAS was determined using LD-clumping [38, 39] based on the LD structure annotated in the 1000 genomes project [40]. SNPs were considered for the clumping analysis if they passed a genome-wide significance threshold of P<5×10^−8^. Following this, SNPs that were within a 500kb region and in LD of at least 0.1 with the top SNPs were included in the clump; SNPs from the region were assigned to the clump if they had an association P<1×10^−5^.

GWAS lookups were performed for the 17 top hits reported in Li et al 2012, Heilmann-Heimbach et al. 2016 and Liu et al. 2016 [13, 15, 16].

Gene-based analyses were performed using MAGMA [41]. SNPs were mapped to genes according to their position in the NCBI 37.3 build map. No additional boundary was added beyond the genes start and stop site. For the autosomal genes the summary statistics from the imputed GWAS were used to derive gene-based statistics using the 1000 genomes (phase 1, release 3) to model linkage disequilibrium. For genes on the X chromosome the genotype data from UK Biobank was used and the gene-based statistic was derived using each participant’s phenotype score.

For the prediction analysis, the GWAS was re-run on a randomly selected cohort of 40,000 individuals, leaving an independent cohort of 12,874 in which to test the polygenic predictor. The methods for the GWAS were identical to those reported for the full sample. The regression weights from the GWAS on the 40,000 cohort were used to construct polygenic scores in the target dataset at P value thresholds of <1×10^−20^, <5×10^−8^, <1×10^−5^, <0.01, <0.05, <0.1, <0.2, <0.5, <1 using PRSice software [42]. PRSice creates polygenic scores by calculating the sum of alleles associated with male pattern baldness across many genetic loci, weighted by their effect sizes estimated from the male pattern baldness GWAS. Thereafter, each threshold was used to discriminate between those with no hair loss and those with severe hair loss via logistic regression with results being reported for the optimal predictor only. A predictor for both the autosomes and X chromosome were built and assessed independently and additively. Receiver operator characteristic (ROC) curves were plotted and areas under the curve (AUC) were calculated using the pROC package in R [43, 44].

SNP-based heritability of baldness was estimated using GCTA-GREML [45] after applying a relatedness cut-off of >0.025 in the generation of the autosomal (but not X chromosome) genetic relationship matrix. Linkage disequilibrium score (LDS) regression analyses [46] were used to generate genetic correlations between baldness and 22 cognitive, anthropometric, and health outcomes, where phenotypic correlations or evidence of shared genetic architecture have been found (**Supplementary Table 6**). Due to the large effects in the *APOE* region, 500kb was removed from around each side of this region and the analysis was repeated. The Alzheimer’s data set without this region is referred to as Alzheimer’s 500kb and makes a total of 23 tests performed and controlled for using False discovery rate (FDR) correction [47]. An overview of the GWAS summary data for the anthropometric and health outcomes is provided in **Appendix 1**.

Partitioned heritability analysis was performed using stratified linkage disequilibrium score (SLDS) regression [48]. SLDS regression examines groups of SNPs sharing the same functional properties and was used to estimate a heritability metric for each group. The goal of this analysis was to determine if a specific group of SNPs made a greater contribution to the total heritability of male pattern baldness than would be expected by the size of the SNP set. Firstly, a baseline model was derived using 52 overlapping, functional categories. Secondly, a cell-specific model was constructed by adding each of the 10 cell-specific functional groups to the baseline model one at a time to the baseline model. Multiple testing was controlled for using FDR correction [47] to the to the baseline model using 52 categories. For the cell-specific analysis the baseline model was first included and the level of enrichment for each cell specific category as derived. Here, 10 tests were controlled for using a FDR correction.

## Acknowledgements

This research was conducted, using the UK Biobank Resource, in The University of Edinburgh Centre for Cognitive Ageing and Cognitive Epidemiology, part of the cross-council Lifelong Health and Wellbeing Initiative (MR/K026992/1). Funding from the Biotechnology and Biological Sciences Research Council (BBSRC) and Medical Research Council (MRC) is gratefully acknowledged. WDH is supported by a grant from Age UK (Disconnected Mind Project).

## Conflicts of Interest

Ian Deary and David Porteous are participants in UK Biobank.

---

**Supplementary Table 1.**
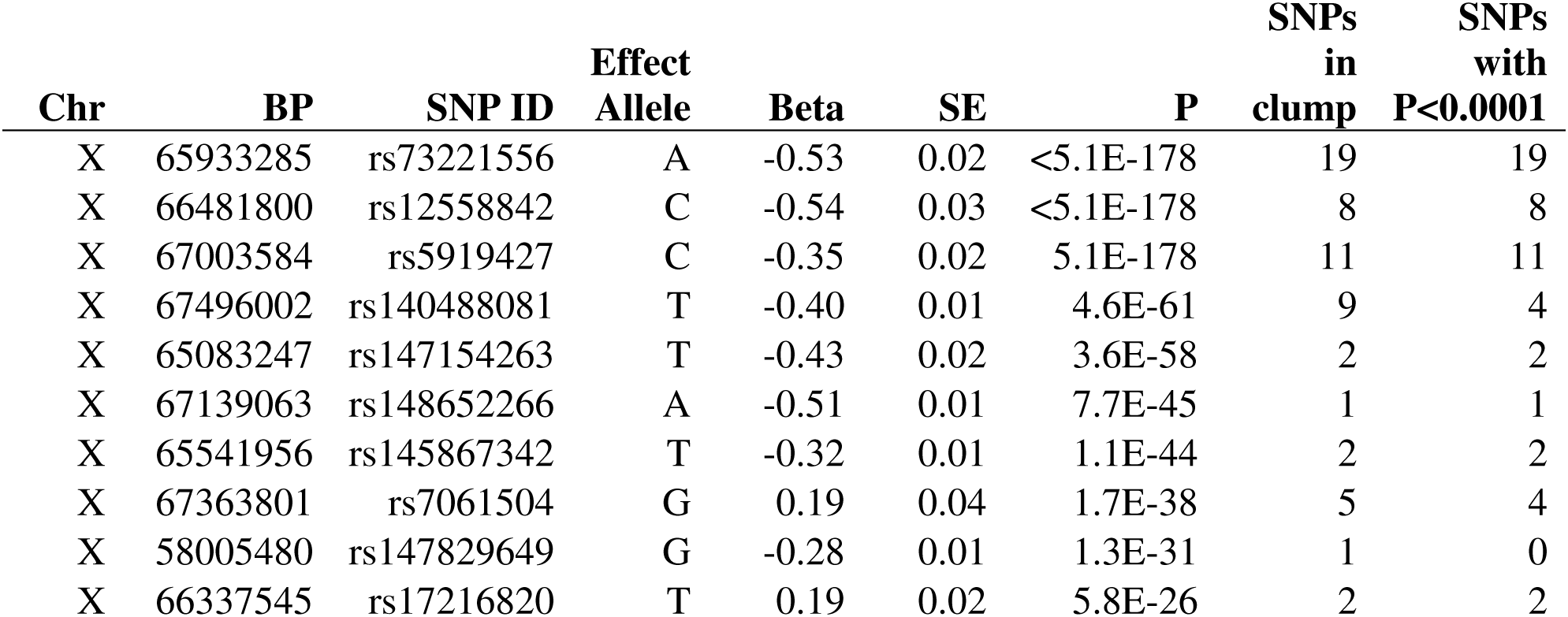
List of top 10 genotyped male-pattern baldness GWAS hits for the X chromosome.

| **Chr** | **BP** | **SNP ID** | **Effect Allele** | **Beta** | **SE** | **P** | **SNPs in clump** | **SNPs with P<0.0001** |
| --- | --- | --- | --- | --- | --- | --- | --- | --- |
| X | 65933285 | rs73221556 | A | -0.53 | 0.02 | <5.1E-178 | 19 | 19 |
| X | 66481800 | rs12558842 | C | -0.54 | 0.03 | <5.1E-178 | 8 | 8 |
| X | 67003584 | rs5919427 | C | -0.35 | 0.02 | 5.1E-178 | 11 | 11 |
| X | 67496002 | rs140488081 | T | -0.40 | 0.01 | 4.6E-61 | 9 | 4 |
| X | 65083247 | rs147154263 | T | -0.43 | 0.02 | 3.6E-58 | 2 | 2 |
| X | 67139063 | rs148652266 | A | -0.51 | 0.01 | 7.7E-45 | 1 | 1 |
| X | 65541956 | rs145867342 | T | -0.32 | 0.01 | 1.1E-44 | 2 | 2 |
| X | 67363801 | rs7061504 | G | 0.19 | 0.04 | 1.7E-38 | 5 | 4 |
| X | 58005480 | rs147829649 | G | -0.28 | 0.01 | 1.3E-31 | 1 | 0 |
| X | 66337545 | rs17216820 | T | 0.19 | 0.02 | 5.8E-26 | 2 | 2 |

**Supplementary Table 2.** Genome-wide significant autosomal gene-based hits (Bonferroni correction of α < 2.769e-06) in the MAGMA gene-based analysis for male pattern baldness. NSNPS is the number of SNPs in the gene.

(See attached Excel spreadsheet)

**Supplementary Table 3.** Genome-wide significant gene-based hits (Bonferroni correction of α < 8.818e-05) in the MAGMA gene-based analysis for male pattern baldness performed on the X chromosome. NSNPS is the number of SNPs in the gene.

(See attached Excel spreadsheet)

**Supplementary Table 4.** Genetic correlations between baldness and the 22 cognitive, health, psychiatric, and anthropometric variables. The heritability Z-score and the mean χ^2^ indicate the level of power to detect association where a heritability Z-score of >4 and a mean χ^2^ >1.02 being considered well powered [46]. None of the 23 tests performed survived FDR control for multiple comparisons.

| Phenotypes | Genetic correlation | Standard error | P-value | Heritability Z-score | Mean $\chi^2$ |
| --- | --- | --- | --- | --- | --- |
| **Cognitive abilities** |  |  |  |  |  |
| Childhood intelligence | -0.039 | 0.064 | 0.544 | 5.830 | 1.076 |
| Educational attainment | -0.046 | 0.038 | 0.221 | 13.250 | 1.226 |
| Verbal-numerical reasoning | -0.034 | 0.048 | 0.482 | 10.526 | 1.168 |
| **Health and Metabolic** |  |  |  |  |  |
| Coronary artery disease | 0.003 | 0.049 | 0.958 | 8.111 | 1.146 |
| HbA1c | -0.033 | 0.054 | 0.541 | 5.231 | 1.060 |
| HDL cholesterol | -0.018 | 0.042 | 0.673 | 5.600 | 1.153 |
| HOMA B | 0.095 | 0.062 | 0.125 | 6.273 | 1.053 |
| HOMA IR | 0.123 | 0.068 | 0.072 | 5.400 | 1.053 |
| LDL cholesterol | -0.043 | 0.047 | 0.361 | 3.654 | 1.141 |
| Obesity | 0.018 | 0.032 | 0.567 | 17.100 | 1.125 |
| Triglycerides | -0.023 | 0.034 | 0.510 | 5.846 | 1.154 |
| Type 2 diabetes | -0.028 | 0.051 | 0.584 | 8.727 | 1.134 |
| Waist to hip ratio | -0.016 | 0.04 | 0.692 | 13.286 | 1.358 |
| **Psychiatric** |  |  |  |  |  |
| ADHD | -0.083 | 0.105 | 0.434 | 2.417 | 1.016 |
| Alzheimer's disease 500bp | 0.058 | 0.078 | 0.462 | 5.545 | 1.106 |
| Alzheimer's disease | 0.064 | 0.096 | 0.506 | 1.909 | 1.115 |
| Autism | -0.035 | 0.052 | 0.498 | 8.519 | 1.059 |
| Bipolar disorder | -0.126 | 0.052 | 0.015 | 10.341 | 1.186 |
| MDD | 0.123 | 0.083 | 0.141 | 5.586 | 1.078 |
| Schizophrenia | -0.055 | 0.03 | 0.068 | 22.208 | 1.814 |
| Neuroticism | 0.091 | 0.078 | 0.241 | 4.500 | 1.057 |
| **Anthropometric/Developmental** |  |  |  |  |  |
| Height | -0.061 | 0.028 | 0.030 | 17.400 | 2.982 |
| Age at menarche | -0.096 | 0.032 | 0.003 | 20.625 | 1.401 |

ADHD, attention deficit hyperactivity disorder; MDD, major depressive disorder

**Supplementary Table 5.** Showing the full output of the partitioned heritability analysis for male pattern baldness. Prop._SNPs refers to the proportion of SNPs from the data set that were a part of the corresponding functional annotation. Statistical significance indicated in bold.

(See attached Excel spreadsheet)

**Supplementary Table 6.**
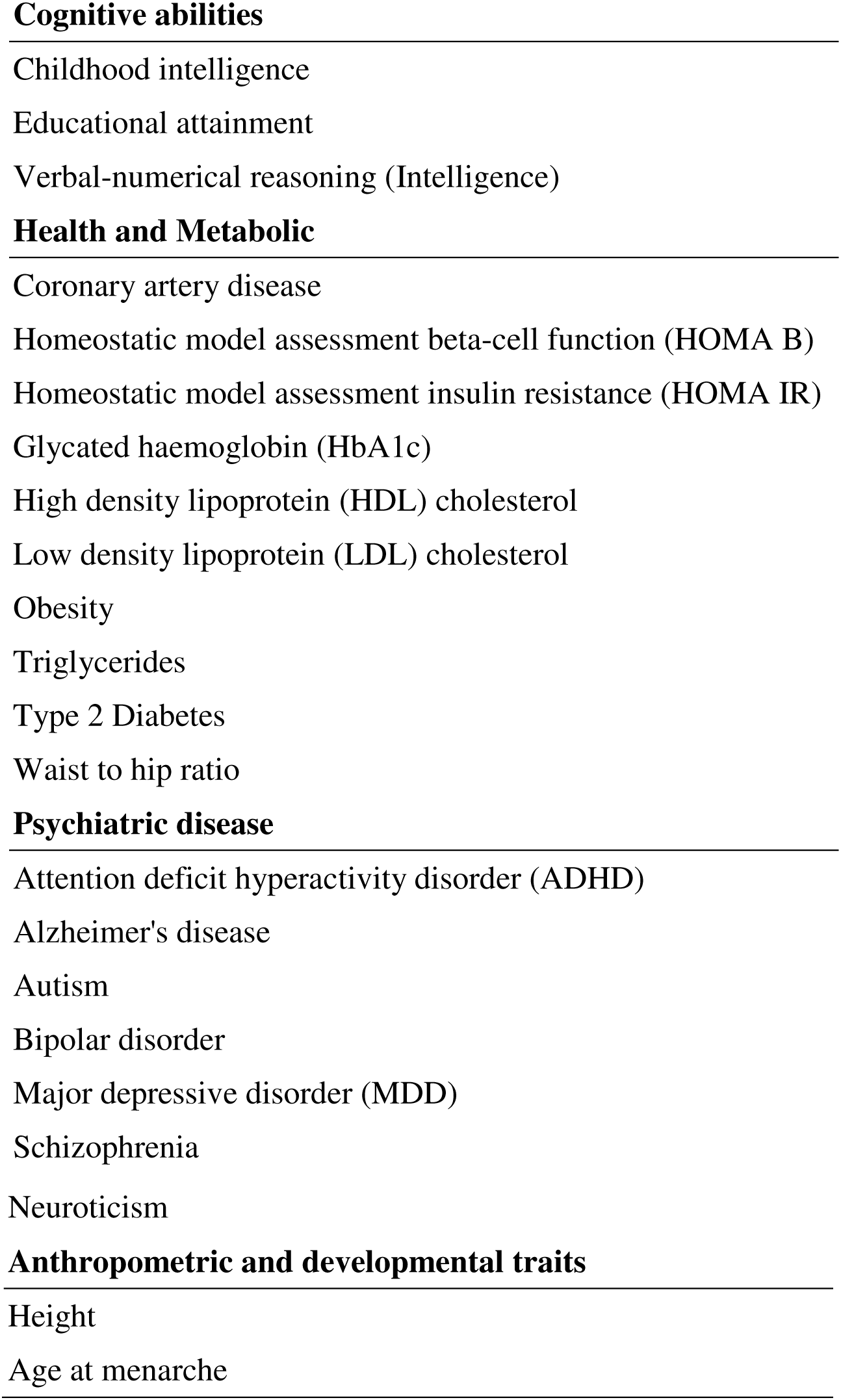
The 22 health-related phenotypes included in the genetic correlation analysis with male pattern baldness. Verbal-numerical reasoning and childhood intelligence were examined as educational attainment (genetic association with baldness reported by Pickrell et al. 2016 [14]) can be used as a proxy phenotype for general cognitive ability. Metabolic traits were included as metabolic disease has been associated with baldness (references noted in the review paper by Heilmann-Heimbach et al. 2016 [16]). Psychiatric disorders were included due to the association between baldness and neurological conditions such as Parkinson’s disease [15]. Genetic correlations have been observed between baldness and the listed anthropometric and developmental traits [14].

## List of Supplementary Figures

**Supplementary Figure 1.**
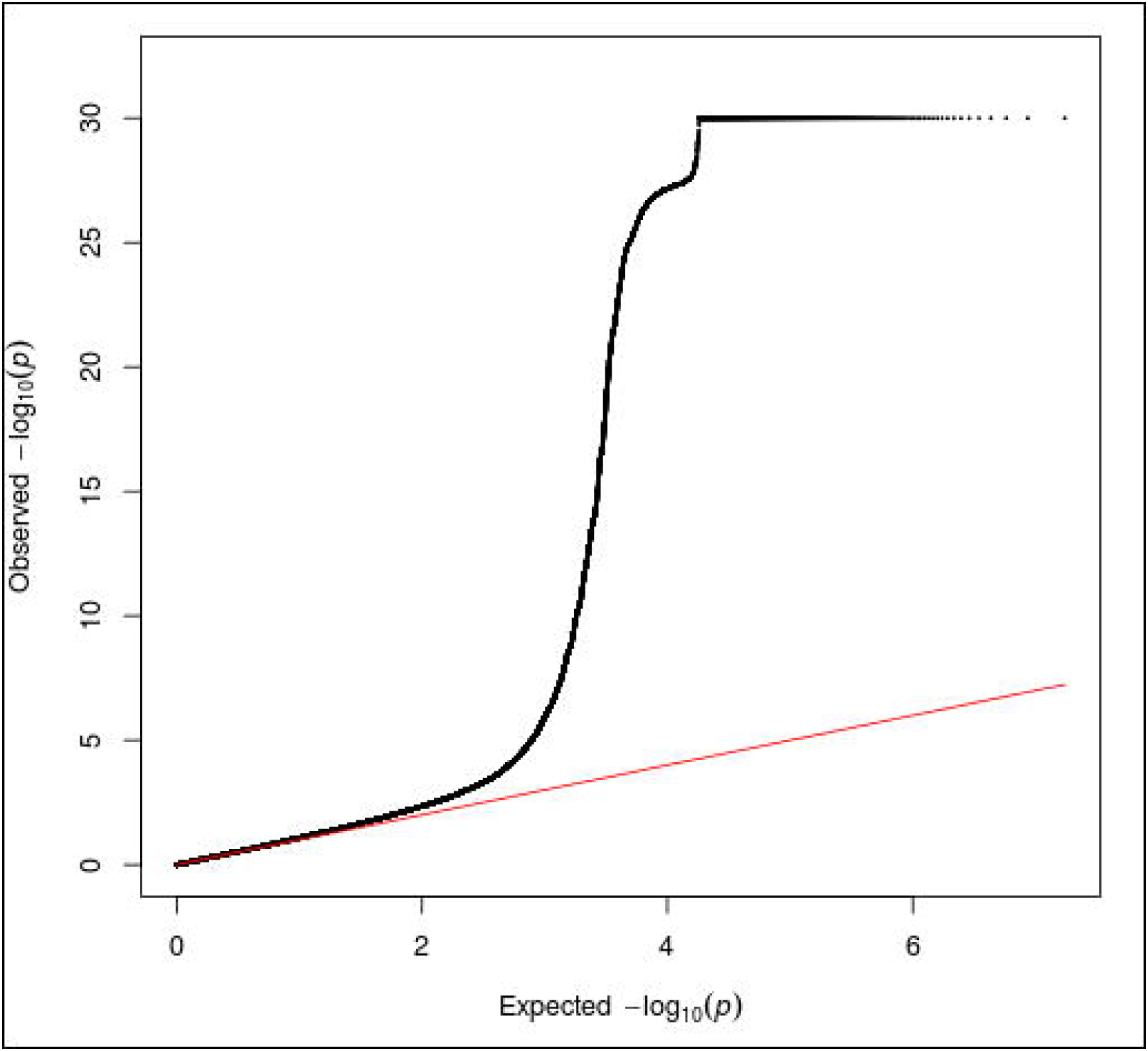
Male-pattern baldness QQ Plot for imputed GWAS of autosomal variants (p-values truncated at 1×10^−30^)

**Supplementary Figure 2.**
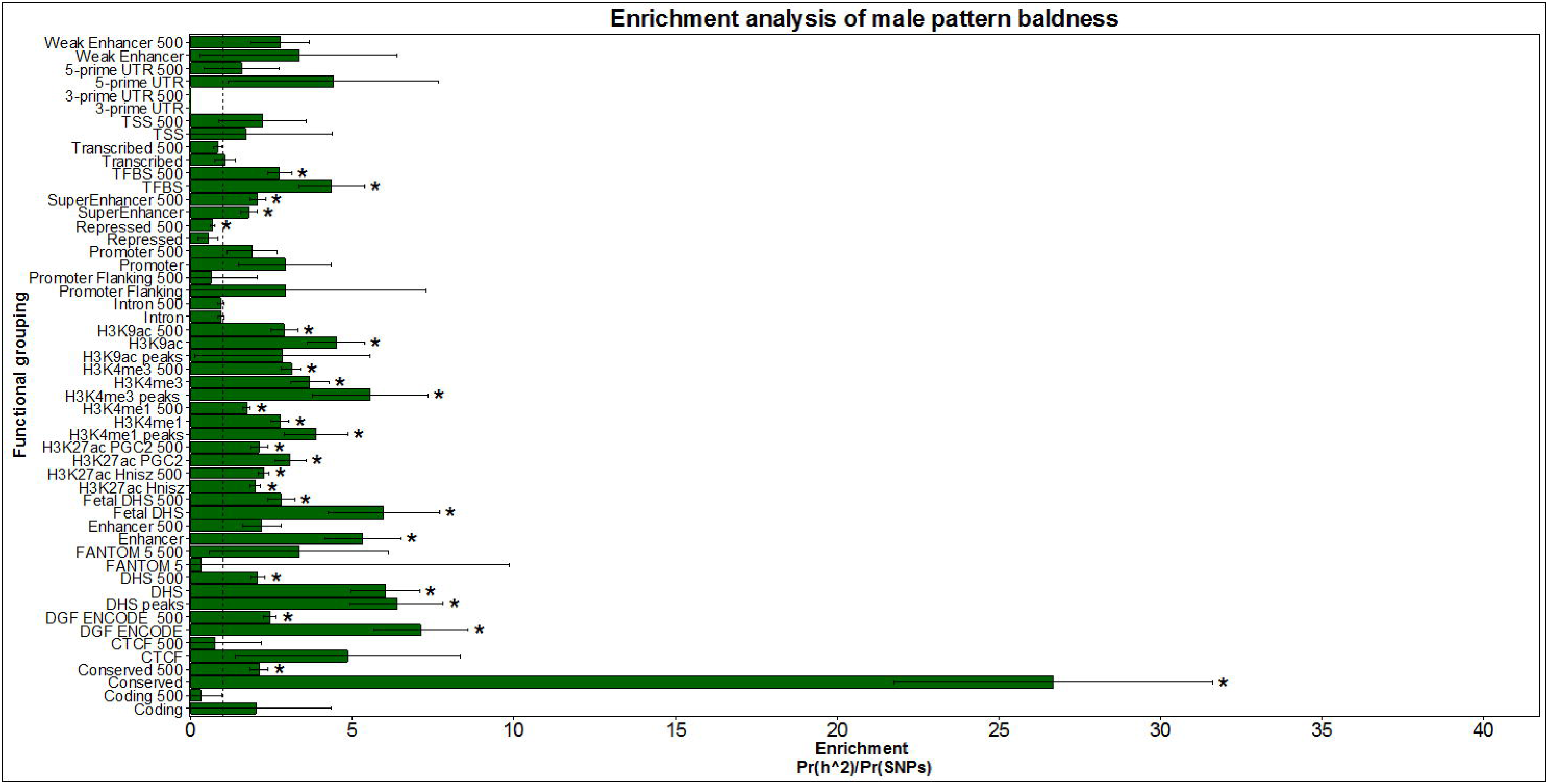
Enrichment analysis for male pattern baldness using the 52 functional categories in 52,874 individuals. The enrichment statistic is the proportion of heritability found in each functional group divided by the proportion of SNPs in each group (Pr(h^2^)/Pr(SNPs)). Error bars are jackknife standard errors around the estimate of enrichment. The dashed line indicates no enrichment found when Pr(h^2^)/Pr(SNPs) = 1. FDR correction indicated significance at P=0.011 indicated by asterisk.

**Supplementary Figure 3.**
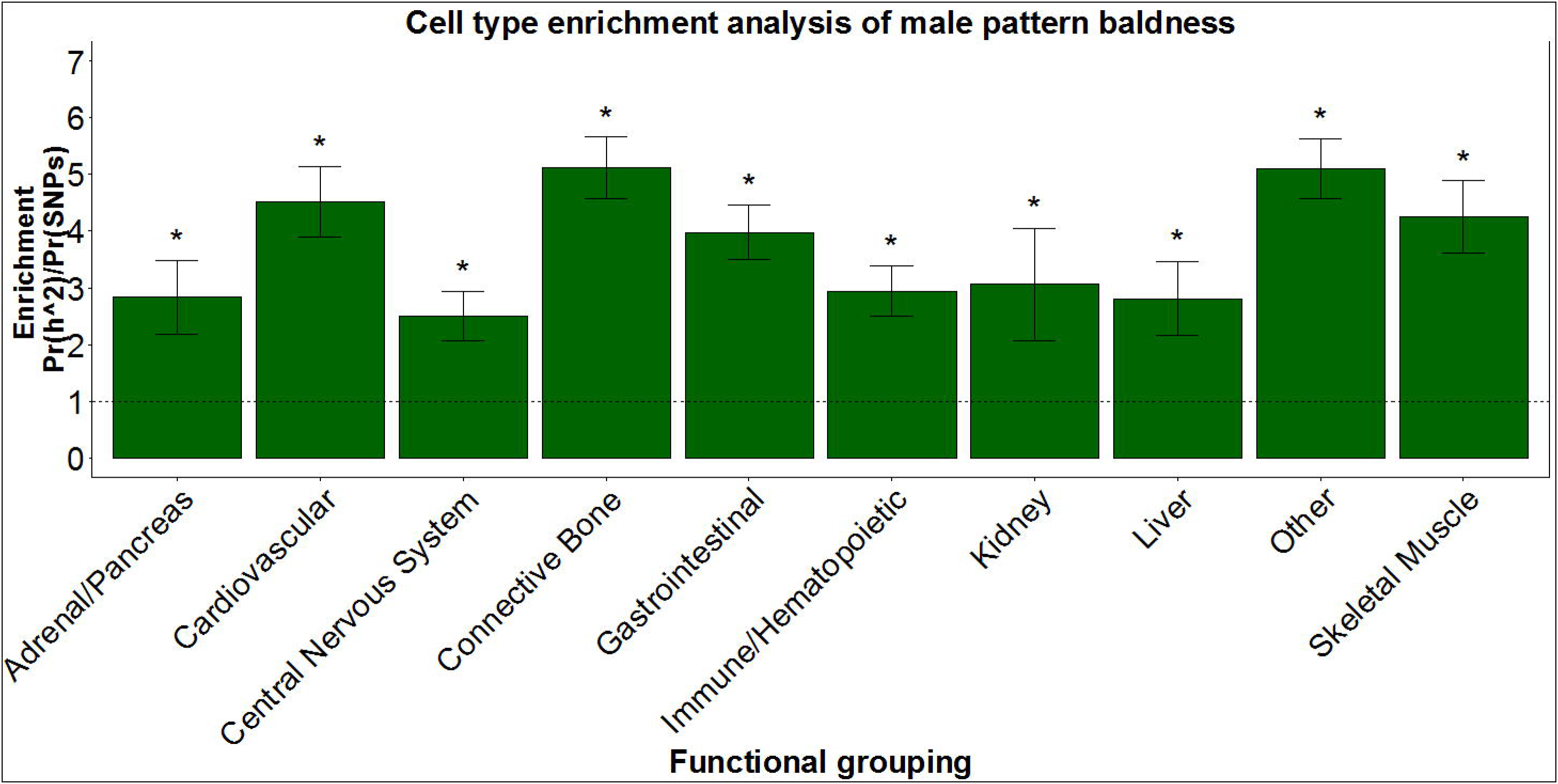
Enrichment analysis for male pattern baldness using the 10 cell specific functional The enrichment statistic is the proportion of heritability found in 52,874 individuals. in each functional group divided by the proportion of SNPs in each group (Pr(h^2^)/Pr(SNPs). Error bars are jackknife standard errors around the estimate of enrichment. The dashed line indicates no enrichment found when Pr(h^2^)/Pr(SNPs) = 1. FDR correction indicated significance at P=0.037 indicated by asterisk.

